# Regulation of Necroptosis and the Type Interferon Response by Monkeypox/mpox Virus

**DOI:** 10.1101/2024.10.31.621358

**Authors:** Jacqueline Williams, James Bonner, Bertram L. Jacobs

## Abstract

Monkeypox/mpox virus (MPXV) has re-emerged as the most important orthopoxvirus infection of humans. Despite being pathogenic, it contains a natural truncation at the N-terminus of its homologue of the vaccinia virus (VACV) innate immune evasion protein, E3, leading to the loss of the first 37 amino acids. Our previous data have shown that VACV E3 protein is required for interferon-resistance of VACV. The N-terminal Z-nucleic acid binding domain is necessary to inhibit induction of necroptosis, as VACV containing an N-terminal deletion (VACV-E3LΔ37N) undergoes rapid ZBP1-dependent necroptotic cell death, through activation of RIPK3, and subsequent phosphorylation and trimerization of the executioner of necroptosis, MLKL. Despite lacking parts of the N-terminus, MPXV has evolved ways in circumventing necroptotic cell death in mouse L929 cells. Our data shows that MPXV infection inhibits phosphorylation of MLKL and does not lead to MLKL trimerization, nor cell death. We show that MPXV inhibits necroptosis in two steps: by degrading RIPK3 through either a caspase-dependent pathway or through the “viral inducer of RIPK3 degradation” (vIRD) and by inhibiting MLKL aggregation. Additionally, MPXV shows interferon sensitivity that is fully ZBP1- and RIPK3- and MLKL-dependent, but independent of necroptosis. Addition of a pancaspase inhibitor (zVAD), or a proteasome inhibitor (MLN4924) does not increase cell death but leads to an increase in interferon sensitivity comparable to VACV-E3LΔ37N. Thus, treatment with an interferon inducer along with a pancaspase or proteasome inhibitor could potentially be a beneficial treatment against MPXV infections.

## Introduction

Monkeypox/mpox virus (MPXV) is an orthopoxvirus that causes smallpox-like disease which has up to a 10% mortality rate, depending on the infectious strain [1]. The global eradication of the smallpox virus has led to the decrease in smallpox vaccinations, which over the years has led to a drastic increase in the number of human MPXV cases [2]. Recently, the Global Commission for the Certification of Smallpox Eradication has named MPXV as the most important orthopoxvirus to infect humans since the eradication of smallpox [3]. In the past MPXV has remained mainly endemic to western and central Africa with recent sporadic cases linking back to travelers who had visited Nigeria. The first MPXV outbreak in the U.S. occurred in 2003 and was traced back to infected rodents imported from Africa [4]. There were also other isolated cases of MPXV spread from 2018-2021 to Israel, the UK, Singapore, and U.S. [5], [6], [7], [8]. In 2022 a worldwide outbreak of mpox occurred with spread throughout the world in people who had not traveled to Africa, with 99,518 worldwide cases in 122 countries [9]. The following year, the cases of MPXV in central Africa started rising and with the emergence of a new MPXV strain, MPXV clade Ib, a more virulent strain, compared to the 2022 MPXV clade IIb strain. Currently, in the Democratic Republic of the Congo (DRC) alone, there are 18,000 suspected MPXV cases, with many cases among children, and 600 MPXV-associated deaths [10]. Two clade Ib MPXV cases have also been reported in Sweden and Thailand in individuals that traveled to Africa [10].

The high rate of transmission of MPXV around the world has become alarming. There are currently two vaccines available that are effective against MPXV: JYNNEOS^TM^ (live, replication incompetent vaccinia virus (VACV)) and ACAM2000^®^ (live, replication competent VACV) [11], however they may be available in limited supplies. According to the Centers for Disease Control and Prevention (CDC), there is no specific treatments specifically against MPXV infections, but there are antiviral drugs that have been shown to be effective against orthopoxviruses. The antiviral ST-246 (also known as Tecovirimat or TPOXX) has been the current drug of choice [11], [12], and brincidofovir and cidofovir are also approved [11]. In a recent trial in Africa, TPOXX did not prove efficacious in people infected with mpox. Given the limited number of antivirals specific against MPXV, development of additional anti-orthopoxvirus drugs would be highly beneficial in combating the current MPXV outbreak. In this paper, we show that an interferon inducer, either with a pancaspase inhibitor or a proteasome inhibitor, could potentially be used as a potential treatment against MPXV infection.

Much of our knowledge of orthopoxvirus pathogenesis and immune evasion comes from the study of VACV. The E3 protein of VACV has been shown to be essential for inhibiting the cell’s antiviral interferon (IFN) response [13], for PKR inhibition [13], [14], [15], [16], [17], and for pathogenesis [18]. The E3 protein is comprised of two conserved domains: the N-terminal Z-form nucleic acid (Z-NA) binding domain (known as ZBD or Z⍺) and the C-terminal double-stranded RNA (dsRNA)-binding domain (dsRBD). Expression of the E3L gene leads to formation of two protein isoforms, one full-length 190-amino acid protein that contains both N- and C-terminal domains (p25), and a smaller protein that is truncated of 37 N-terminal amino acids (p20), which is the result of leaky scanning past the first AUG start codon (p20). A genomic comparison between MPXV and VACV showed that the E3 homologue of MPXV, F3, is 92% identical at the nucleotide level and 88% at the protein level to the VACV E3 protein. The F3 protein retains a complete, fully functional dsRBD, however, the first 37 amino acids of the N-terminal ZBD are missing, resembling the smaller p20 VACV E3 protein [19].

Apoptosis and necroptosis are linked death-receptor activated pathways that result in cell death. In apoptotic conditions, active caspase-8 will cleave and inactivate receptor interacting protein kinase 1 (RIPK1) and RIPK3 leading to apoptosis [20]. When caspase 8 is inhibited, RIPK1 and RIPK3 form a complex leading to the phosphorylation and oligomerization of mixed lineage kinase like-domain protein (MLKL), the final executioner of MLKL [21], [22], [23], [24]. Phosphorylated MLKL will translocate to the plasma membrane where it trimerizes and permeabilizes the membrane, resulting in cell death [25], [26]. RIPK1 and RIPK3 each contain a receptor-interacting protein homotypic interaction motif (RHIM) domain that can mediate protein-protein interaction. In addition, a RHIM is present in the TLR-adaptor protein TRIF, as well as Z-DNA binding protein 1 (ZBP1, also known as DAI or DLM1) [27].

We have previously shown that the VACV N-terminus of E3 protein is necessary to inhibit IFN-primed necroptosis and for IFN resistance in L929 cells. When L929 cells were pretreated with IFN⍺ and infected with VACV, mutants that were missing part or the entire Z-NA binding domain of E3 (VACV-E3LΔ37N or VACV-E3LΔ83N), showed an IFN sensitive phenotype, compared to wild-type VACV (wtVACV) that was fully IFN resistant. Infection with these VACV mutants also led to rapid necroptotic cell death, necroptosis, that was ZBP1- and RIPK3-dependent [28]. Here, we explore how MPXV responds to the host antiviral IFN pathway and ways it may have evolved to remain pathogenic despite having a truncation in its E3 homologue.

## Results

### MPXV is IFN sensitive and does not induce cell death

Since the E3 homologue in MPXV has a natural truncation at the N-terminus similar to VACV-E3LΔ37N, we wanted to see if MPXV was IFN-sensitive in L929 cells, similar to what our lab had previously observed with VACV N-terminal mutants, VACV-E3LΔ37N and VACV-E3LΔ83N [28], which are missing all or part of the N-terminal Z-NA binding domain of E3 (Fig. 1A). L929 cells were pretreated with increasing doses of mouse IFN⍺ for 18 hours to induce expression of ZBP1 and infected with 100 plaque forming units (PFU) of wtVACV, VACV-E3LΔ37N, or MPXV. As shown in Fig. 1B, wtVACV is fully IFN resistant, VACV-E3LΔ37N is IFN sensitive, and MPXV is IFN sensitive with an intermediate phenotype between wtVACV and VACV-E3LΔ37N. These data show that in L929 cells, IFN⍺ can reduce plaque formation of MPXV with 10 U of IFN reducing plaques by about 50%.

**Figure 1.**
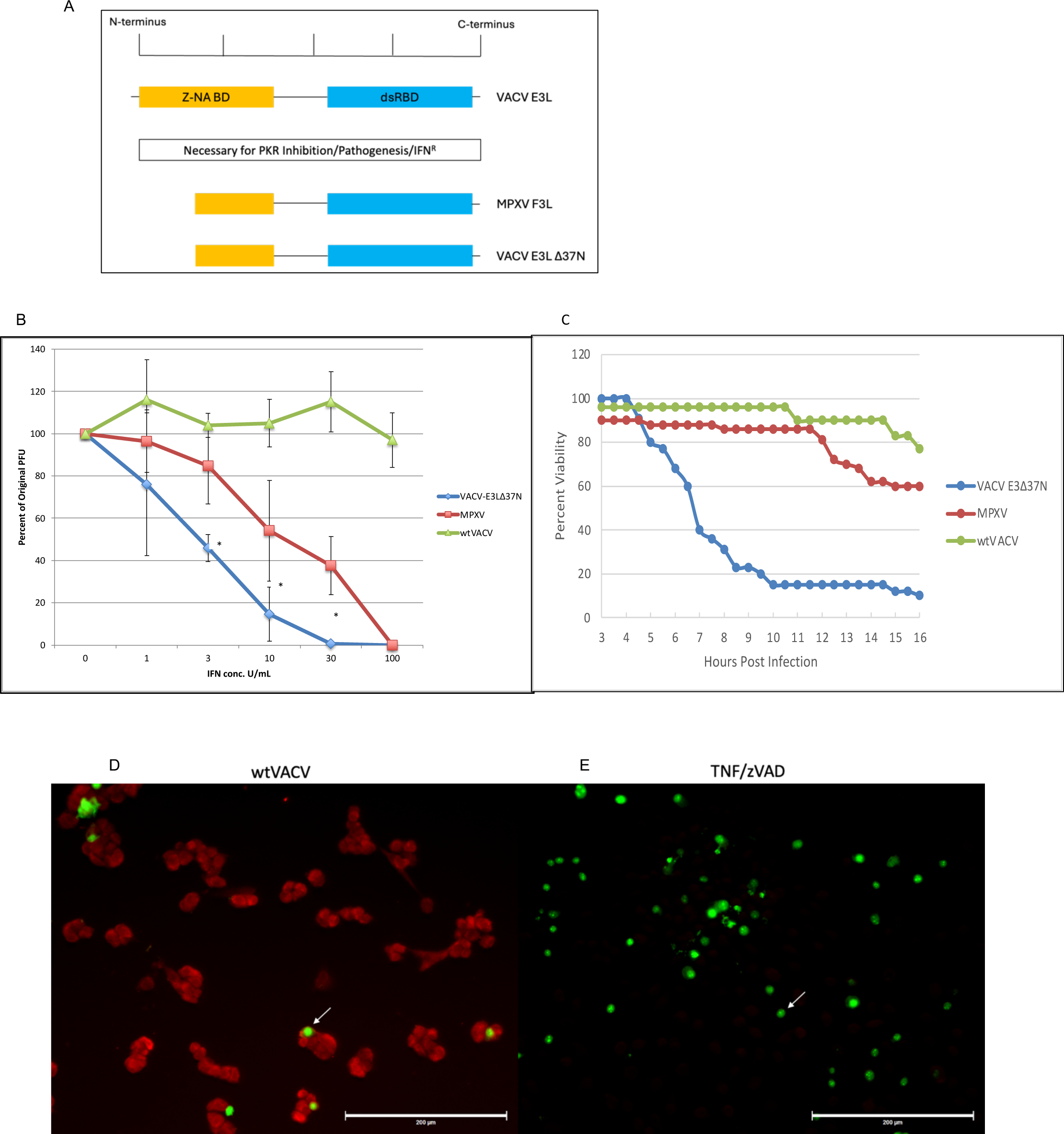
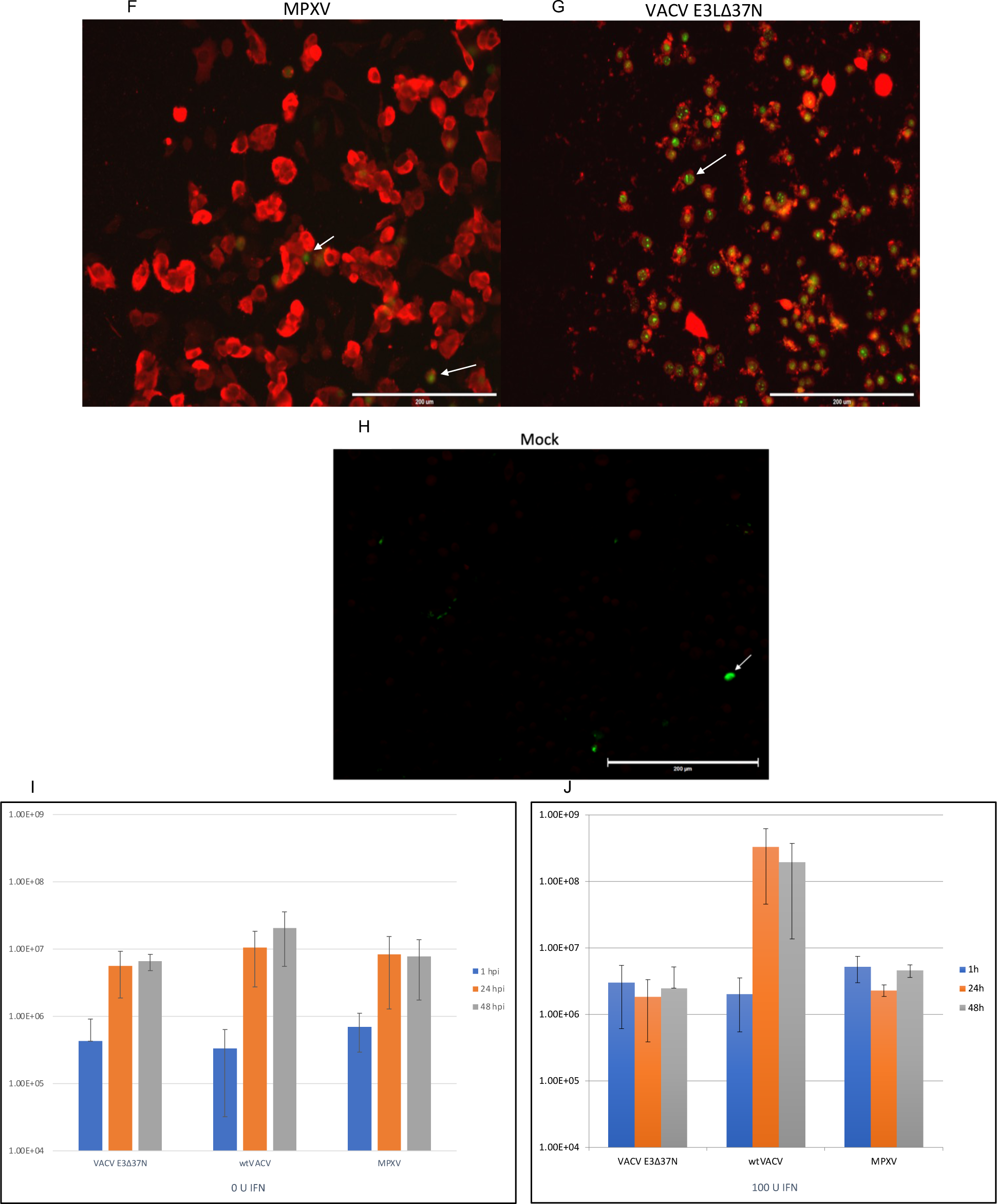
MPXV is IFN sensitive and does not induce cell death. A) E3 schematic. B) IFN plaque reduction assay showing MPXV has an intermediate IFN phenotype between wtVACV and VACV-E3Δ37N. C) SYTOX green cell viability assay showing VACV-E3Δ37N causes rapid cell death while MPXV and wtVACV do not. D-H) IF with LIVE/DEAD green fixable stain to visualize infected versus dead cells. Cells were mock treated, treated with TNF/zVAD, or infected with wtVACV, MPXV, and VACV-E3Δ37N. The red is stained for E3 and the green is dead cells. The white arrow shows dead cells. I) 1-step viral growth curves with wtVACV, VACV-E3Δ37N, and MPXV at an MOI of 5 and harvested at 1, 24, and 48 hours post infection and plaqued in BSC40 cells. J) Same as I, except cells were pretreated with 100 U of IFN⍺ for 18 hours. *p=<.05 when comparing VACV-E3Δ37N and MPXV.

IFN-sensitivity of VACV-E3LΔ37N and VACV-E3LΔ83N is associated with ZBP1-dependent necroptotic cell death [28], [29]. The ability of MPXV to induce cell death was examined using live imaging in IFN-primed L929 cells. The cells were pretreated with 100 U of mouse IFN⍺ for 18 hours and infected with VACV-E3LΔ37N or MPXV at an MOI of 5 and the cells were visualized every hour. Cell viability was examined by examining plasma membrane integrity using a SYTOX exclusion assay. Despite having a natural truncation in the N-terminus of the E3 homologue, MPXV infection did not lead to rapid cell death like we see with VACV-E3LΔ37N, instead they remained closer to wtVACV levels (Fig. 1C). Using IF, we next looked at cell death with a fixable cell death stain and E3 protein expression. As shown in Fig. 1D, cells infected with wtVACV stained red for E3 and a few dead cells stained green for cell death. Cells that were treated with TNF/zVAD all stained green (Fig. 1E), while mock had no E3 staining and very few dead cells (Fig. 1H). The majority of the L929 cells infected with MPXV stained positive for E3 while very few cells stained green for dead cells, compared to VACV-E3LΔ37N (Fig. 1F and G, respectively). This data demonstrates that the lack of cell death is not due to absence of virus infection. This also indicates that MPXV has evolved mechanisms of immune evasion independent of the E3 homologue.

We have also evaluated MPXV replication in L929 cells under single cycle replication conditions, in the presence and absence of IFN. A viral growth curve was performed in which the cells were pre-treated with 100 units of IFN for 18 hours and infected with VACV-E3LΔ37N, wtVACV, or MPXV at an MOI of 5 and the samples were harvested at 1, 24, and 48 hours post infection. The viral titers were then determined by plaquing in BSC40 cells. In the absence of IFN, VACV-E3LΔ37N and MPXV titers increased by 1-log and wtVACV titers increased 1.5-logs (Fig. 1I). In the presence of IFN, both VACV-E3LΔ37N and MPXV failed to replicate while wtVACV titers increased 2-logs (Fig. 1J). This data shows that IFN⍺ can efficiently inhibit MPXV viral replication *in vitro* under single cycle conditions.

### IFN sensitivity is dependent on ZBP1, RIPK3, and MLKL but not PKR

IFN-sensitivity of E3LΔ37N and VACV-E3LΔ83N is dependent on the ZBP1/RIPK3/MLKL pathway. We wanted to ask if IFN sensitivity of MPXV was dependent on ZBP1, RIPK3, and MLKL, despite not leading to necroptotic cell death. CRISPR/Cas9 L929 cells that failed to make these proteins (Fig. 2A) were seeded and pretreated with increasing doses of IFN⍺ (0-100 U/mL) for 18 hours. IFN sensitivity was examined by infecting CRISPR KO L929 cells with 100 PFU of wtVACV, VACV-E3LΔ37N, VACV-E3LΔ26C, or MPXV. All viruses were IFN resistant upon deletion of ZBP1, and replicated efficiently in the presence of IFN, except for VACV-E3LΔ26C (Fig. 2B). VACV-E3LΔ26C has 26 amino acids deleted from the C-terminus of E3, which is the dsRNA-binding domain and is required for inhibition of IFN-inducible dsRNA-sensing pathways, PKR and OAS and IFN resistance, but not for inhibition of necroptosis [14], [28], [30]. Similar results were seen in RIPK3 knock-out L929 cells (Fig. 2C). In the L929 MLKL KOs, the IFN sensitivity of MPXV is completely dependent on MLKL. MPXV IFN resistance increased to wtVACV levels but the VACV N-terminal mutant, VACV-E3LΔ37N, increased to 100% resistance with 1 U of IFN and decreased slightly to give 50% resistance at 100 U of IFN. This could be due to ZBP1 and RIPK3 acting in another pathway beyond necroptosis, such as NFkB, apoptosis, or inflammation. VACV-E3LΔ26C remained IFN sensitive, as expected (Fig. 2D). We tested PKR knock-out cells as a control since induction of necroptosis in L929 cells by VACV-E3LΔ37N and VACV-E3LΔ83N has been shown to be PKR-independent [29]. When PKR was knocked out of L929 cells, IFN treatment was able to limit viral replication of VACV-E3LΔ37N and to an intermediate level MPXV, similar to WT L929 cells, despite PKR being absent (Fig. 2E). This demonstrates PKR is dispensable for IFN sensitivity of MPXV in L929 cells. Next, we tested the IFN sensitivity of the highly pathogenic clade I MPXV Zaire strain in our L929 necroptotic KO cell lines. As shown in Fig. 2F, MPXV Zaire was IFN sensitive in WT L929 and in the PKR KOs, but fully IFN resistant in the ZBP1, RIPK3, and MLKL KO cells, similar to the clade II MPXV 7-61 strain.

**Figure 2.**
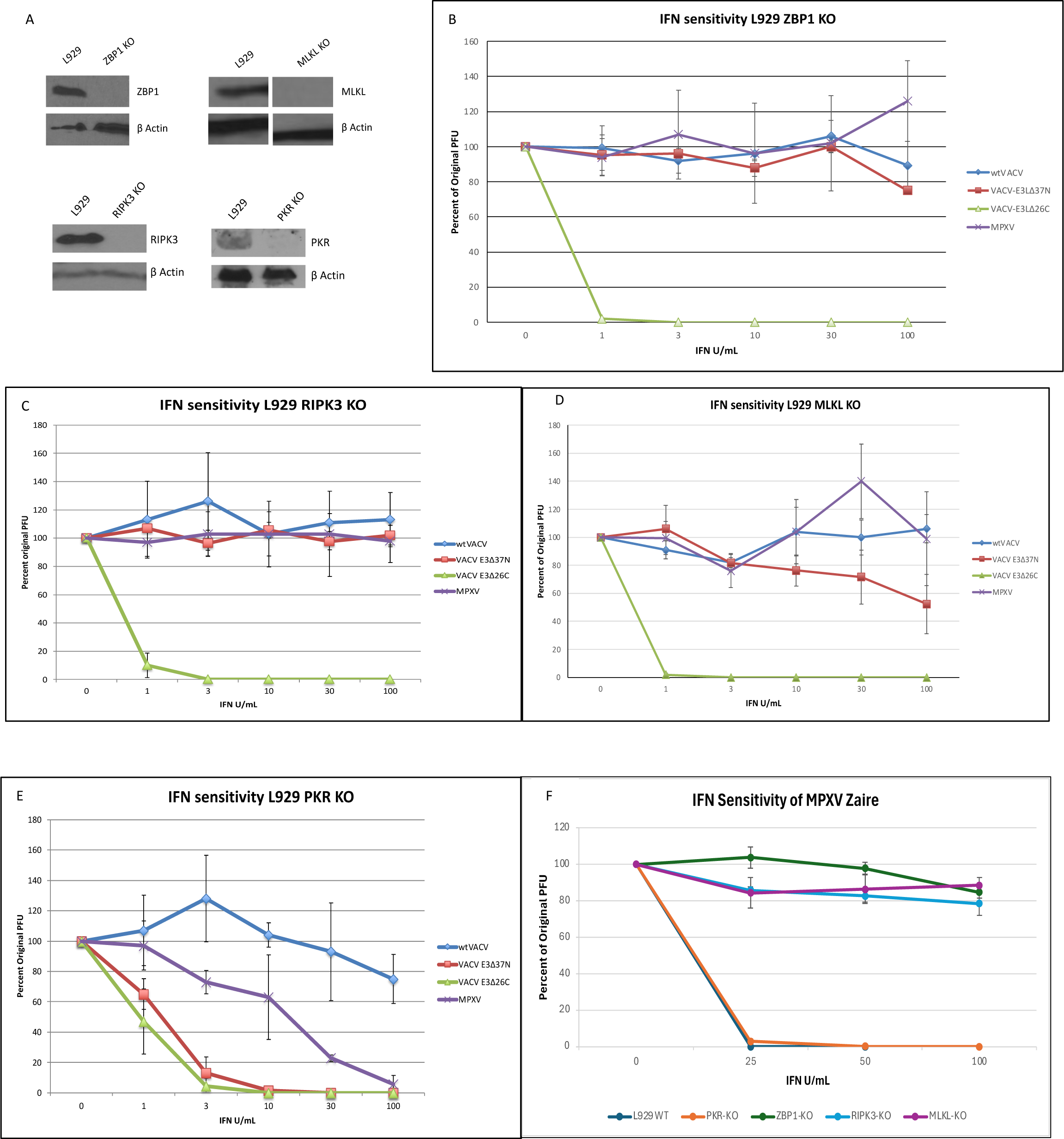
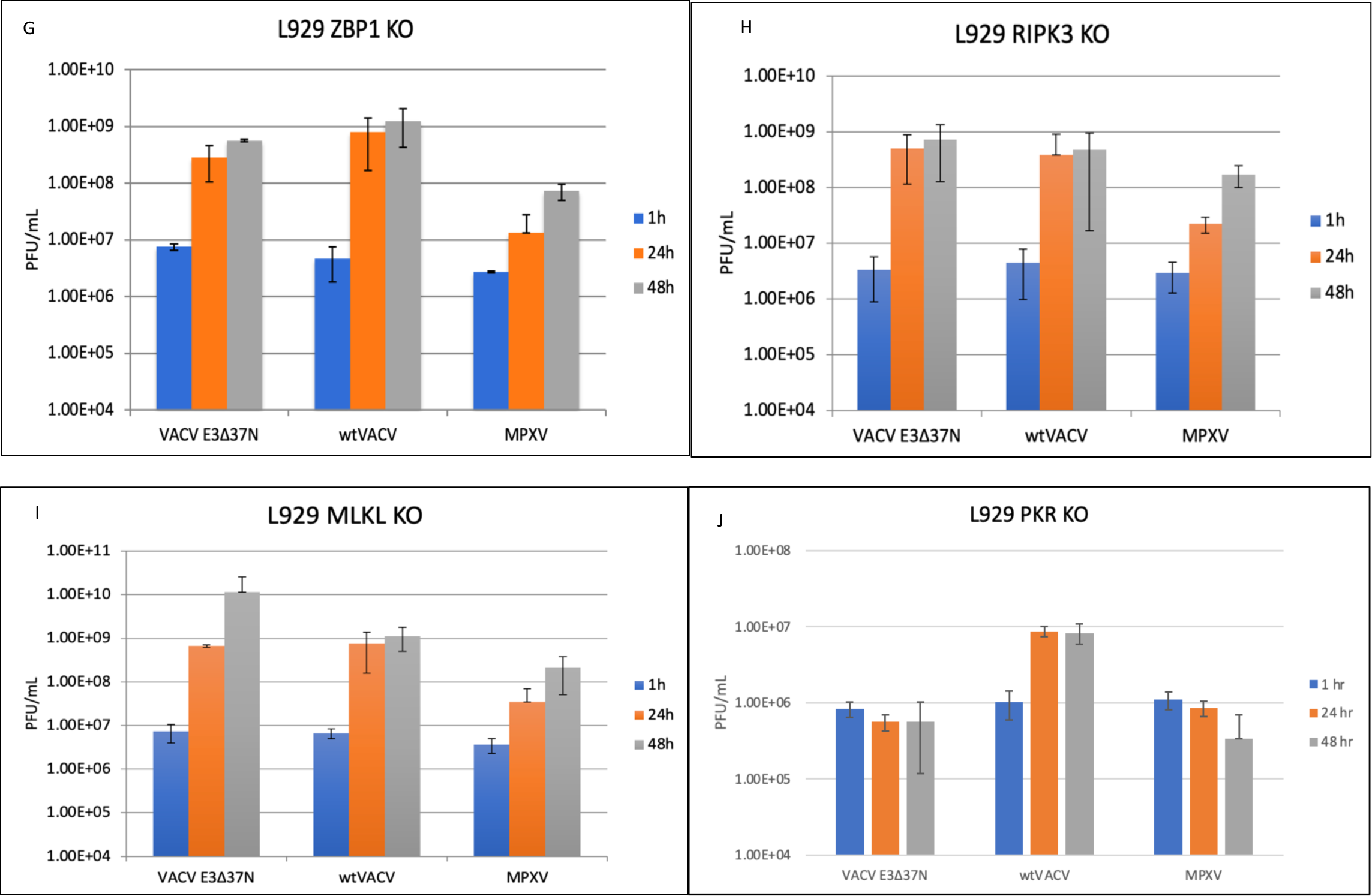
IFN sensitivity is dependent on ZPB1, RIPK3, and MLKL but not PKR. A) Western blot confirming CRISPR KO L929 cells. B-E) IFN plaque reduction assays with CRISPR ZBP1, RIPK3, MLKL, and PKR KO L929 cells. F) IFN plaque reduction assay in WT L929, PKR-KO, ZBP1-KO, RIPK3-KO, and MLKL-KO with MPXV Zaire. G-J) 1-step viral growth curves in ZBP1, RIPK3, MLKL, and PKR KO L929 cells. Cells were pretreated with 100 U IFN⍺ and infected with wtVACV, VACV-E3Δ37N, or MPXV at an MOI of 5. Viral titers were determined by plaquing in BSC40 cells.

We confirmed that ZBP1, RIPK3, and MLKL is required for IFN resistance by looking at viral growth in a 1-step growth curve. The KO cells were pretreated with IFN and infected with VACV-E3LΔ37N, wtVACV, or MPXV at an MOI of 5. The cells were harvested at the indicated time points and the viruses were plaqued in BSC40s to determine the viral titers. VACV-E3LΔ37N replicated 1.5-logs in ZBP1 KO, 2-logs in RIPK3 KO, and 3-logs in MLKL KO cells. wtVACV replicated 2-logs in all the KO cells, and MPXV replicated 1.5-logs in ZBP1 and MLKL KO cells and 2-logs in RIPK3 KO cells (Fig. 2G-I). In PKR KO cells, VACV-E3LΔ37N and MPXV failed to replicate while wtVACV replicated 1-log (Fig. 2J). Thus, MPXV and VACV-E3LΔ37N viral replication in the presence of IFN is dependent ZBP1, RIPK3, and MLKL, but not PKR.

### MPXV vIRD triggers RIPK3 degradation and inhibition of necroptosis

We have previously shown that the E3 protein of VACV is necessary to inhibit viral-induced ZBP1-dependent necroptosis, as the E3 N-terminus deletion mutants caused rapid cell death due to phosphorylation and trimerization of MLKL in L929 cells [28]. We were interested in determining if MLKL becomes phosphorylated and trimerizes in cells infected with MPXV, because MPXV contains a truncated N-terminal E3 homologue, similar to the VACV-E3LΔ37N construct. Thus, we expected that MPXV would likewise trigger MLKL trimerization and necroptosis. L929 cells were infected with wtVACV, MPXV, or VACV-E3LΔ37N, and lysates were harvested at 2, 4, 6, and 8 hours post-infection, immunoblotted, and probed for phosphorylated MLKL. Phosphorylated MLKL was detectable as early as 4 hours post infection with VACV-E3LΔ37N, but not in MPXV or wtVACV infected cells (Fig. 3A), indicating a lack of necroptosis in MPXV infected cells. Since necroptosis is a highly inflammatory process, cells have evolved ways to inhibit this pathway when not necessary. Caspase-8 has been shown to inhibit necroptosis by cleaving RIPK3 and deletion of caspase-8 is lethal in mice embryos which can be rescued by deletion of RIPK3 or MLKL [20], [31],[32], [33], [34]. Inhibition of necroptosis has also been shown to occur with cowpox virus (CPXV) infections through its “viral inducer of RIPK3 degradation” (vIRD). The vIRD contains six ankyrin repeats at the N-terminus and an F-box at the C-terminus. It was shown to bind the host SKP1-Cullin1-F-box (SCF) machinery to target RIPK3 for proteasomal degradation [35]. We wanted to ask if the reason MPXV was not inducing necroptosis was because RIPK3 is being degraded. As shown in figure 3B, cellular expression of RIPK3 was unaffected by infection with wtVACV or VACV-E3LΔ37N, as expected, likely because VACV vIRD is truncated and lacks the C-terminal F-Box [35]. Conversely, RIPK3 expression was lower in MPXV-infected cells, than in VACV-infected cells, and RIPK3 was completely degraded in CPXV-infected cells. We pretreated L929 cells with zVAD-FMK to see if inhibition of caspases would restore RIPK3 levels. In wtVACV and VACV-E3LΔ37N infected cells RIPK3 levels were present in high amounts and addition of zVAD restored RIPK3 levels in MPXV and CPXV infected cells (Fig. 3C). We detected phosphorylation of MLKL at 6 and 8 hours in MPXV infected cells with the addition of zVAD (Fig. 4A), but this did not lead to trimerization of MLKL (Fig. 3F) or necroptotic cell death. This shows that MPXV reduces RIPK3 levels to inhibit MLKL phosphorylation.

**Figure 3.**
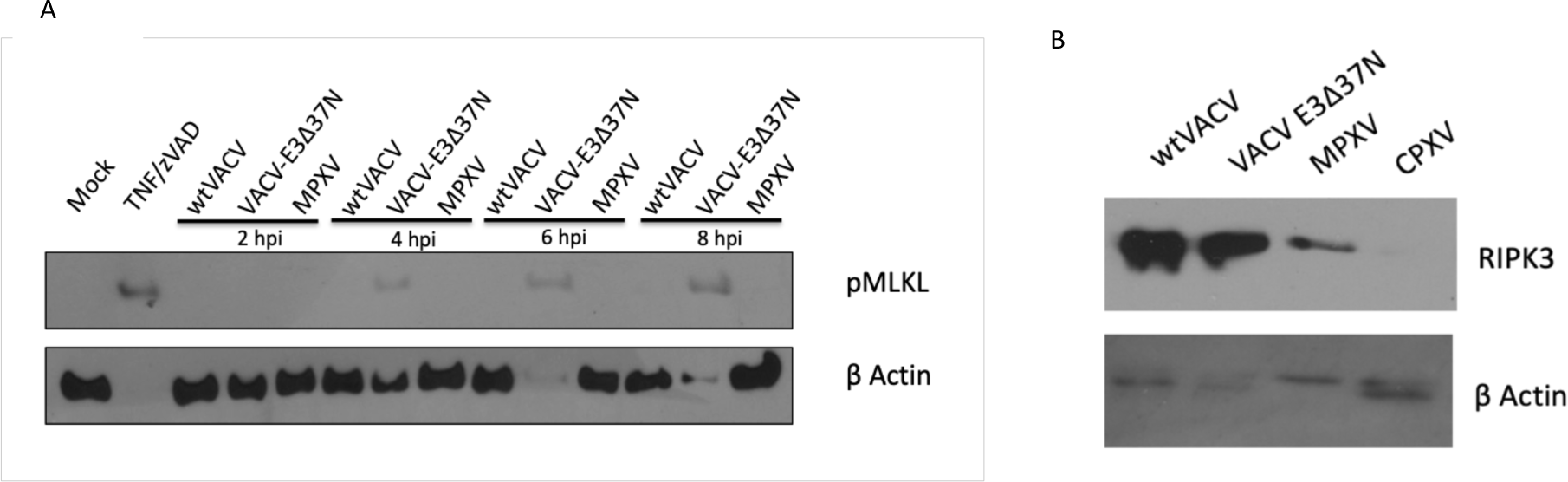
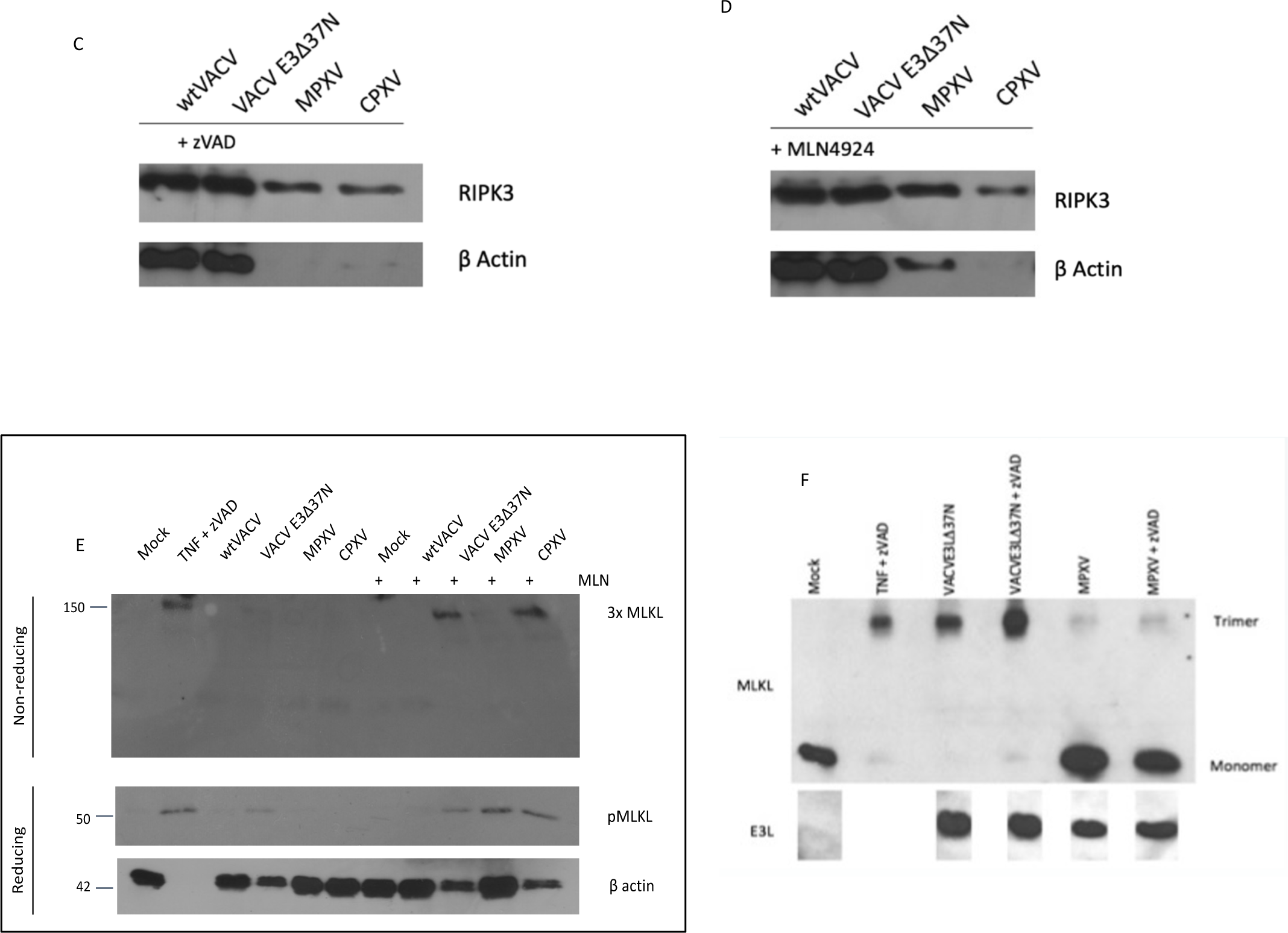
MPXV vIRD leads to RIPK3 cleavage and necroptosis inhibition. A) Western blot showing a time course of phosphorylated MLKL. Viruses infected at an MOI of 5 and L929 cells were pretreated with 100 U IFN⍺. B) Western blot showing RIPK3 degradation in MPXV and CPXV infected cells. C) Western blot for RIPK3 degradation with zVAD. D) Western blot for RIPK3 degradation with MLN4924. E) Western blot for phosphorylation and trimerization of MLKL with and without MLN4924. F) Western blot in non-reducing conditions for trimerized MLKL.

**Figure 4.**
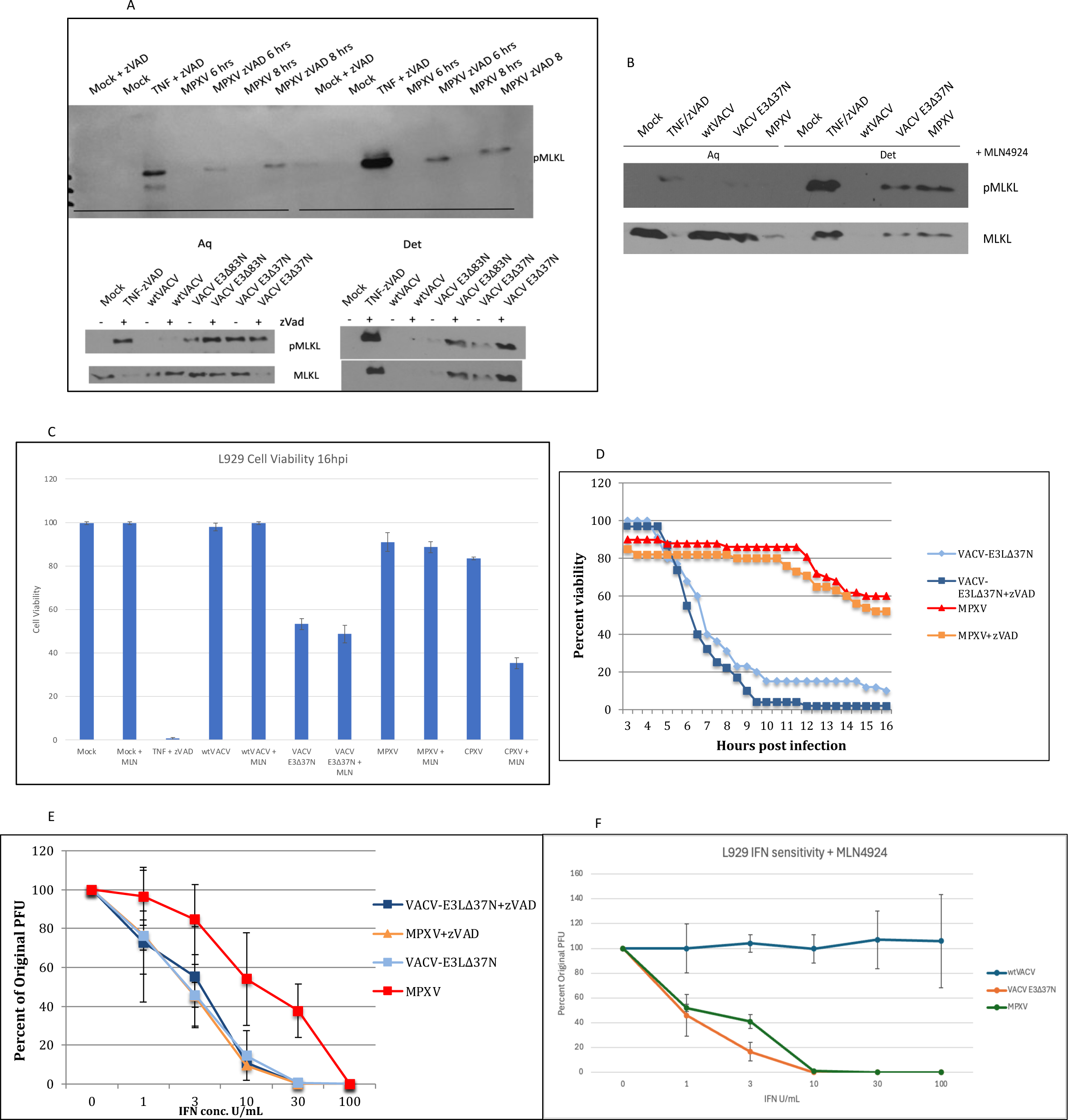
Caspase and neddylation inhibition leads to increased IFN sensitivity and MLKL membrane localization, but does not lead to necroptosis. A) Western blot showing phosphorylation of MLKL localizing to membranes in the presence of zVAD. Viral samples were lysed with TX-114 allowing for separation. B) Same as A, but with MLN4924 instead of zVAD. C) L929 cell viability assay with and without neddylation inhibitor, MLN4924. D) SYTOX green L929 cell viability assay with the addition of pancaspase inhibitor, zVAD. E) L929 IFN plaque reduction assay with VACV-E3Δ37N and MPXV in the presence of zVAD. F) L929 IFN plaque reduction assay with MLN4924.

Next, we used the neddylation inhibitor, MLN4924 (ApexBio), to inhibit RIPK3 degradation to attempt to induce necroptosis in MPXV and CPXV infected cells [35]. L929 cells were pretreated with MLN4924 and infected with the indicated viruses (MOI 5) for 16 hours and RIPK3 levels were analyzed via western blot. wtVACV and VACV-E3LΔ37N expressed RIPK3 to high levels and addition of MLN4924 restored RIPK3 levels in MPXV and CPXV infected cells (Fig. 3D). We confirmed necroptosis by performing a western blot and probing for phosphorylated and trimerized MLKL. Cells infected with CPXV showed phosphorylated and trimerized MLKL only with the addition of MLN4924. Cells infected with wtVACV had no phosphorylated or trimerized MLKL and addition of MLN4924 to VACV-E3LΔ37N infected cells gave a stronger signal for phosphorylated and trimerized MLKL. Addition of MLN4924 to MPXV infected cells enhanced phosphorylation but not trimerization of MLKL (Fig. 3E). This data suggests that MPXV vIRD triggers RIPK3 degradation and inhibits MLKL phosphorylation and trimerization.

### Caspase and neddylation inhibition of MPXV leads to MLKL membrane localization and increased IFN sensitivity without necroptosis

Although zVAD and MLN4924 prevent RIPK3 degradation and potentiate MPXV-induced MLKL phosphorylation, MLKL oligomerization remains unseen. We next sought to determine the next downstream step in the necroptotic pathway where MLKL activation was inhibited by MPXV. We infected L929 cells with the indicated viruses and then separated cytosolic and membrane-bound proteins by using the non-ionic detergent Triton X-114 (TX114) in our lysis buffer. When heated above 22℃, solutions of TX114 resolve into aqueous (Aq) and micellar detergent (Det) phases that can be centrifugally separated. The Aq samples will then contain hydrophilic solutes while the Det samples will contain hydrophobic solutes like membrane-bound proteins [36], allowing us to follow phosphorylated MLKL from the cytoplasm to the membrane in MPXV infected cells. In lysates from TNF/zVAD-treated cells, phosphorylated MLKL was detected in both the Aq and Det samples with a stronger signal in the Det samples. Phosphorylated MLKL was not detected in either the Aq or Det extract from wtVACV-infected cells, consistent with our data that wtVACV does not induce necroptosis. In VACV N-terminal deletion mutants, phosphorylated MLKL is detected in the Aq and Det extracts from cells that were infected with VACV N-terminal mutants, a signal that was enhanced by zVAD. In MPXV infected cells, phosphorylated MLKL is not detected in the Aq or Det phase, but it is clearly visible in both fractions with the addition of zVAD starting at 6 hours post infection (Fig. 4A). Addition of MLN4924 gave similar results showing phosphorylated MLKL localizing to the membrane in MPXV and VACV-E3LΔ37N infected cells (Fig. 4B). Next, necroptosis was investigated by infecting with the indicated viruses (MOI 5) with or without MLN4924 and cell death was measured at 16 hours post infection. Mock and MLN4924 treated cells showed no toxicity, TNF/zVAD treated cells underwent necroptosis, wtVACV and wtVACV with the addition of MLN4924 did not induce cell death in wtVACV-infected cells, while VACV-E3LΔ37N induced necroptosis that was unaffected by MLN4924. Interestingly, MLN4924 potentiated cytotoxicity in CPXV-infected cells but not in MPXV-infected cells (Fig. 4C). In summary, MPXV triggers the phosphorylation and membrane-localization of MLKL only in the presence of the pancaspase inhibitor zVAD or the neddylation inhibitor MLN4924. Finally, even when these inhibitors are present and MLKL phosphorylation is restored, MLKL oligomerization and necroptosis does not occur within MPXV-infected cells.

The default death pathway for virally infected cells is apoptosis. When caspase-8 is inhibited, RIPK3 can be activated through RHIM interactions with RIPK1, ZBP1, or TLR3/TLR4 (TRIF) and lead to MLKL activation and cell death. We wanted to see if the pancaspase inhibitor, zVAD-FMK could induce necroptosis in MPXV infected cells. The addition of zVAD had no significant effect on cell survival in MPXV or VACV-E3Δ37N infected cells, as measured by SYTOX green exclusion. Despite inhibiting RIPK3 cleavage in MPXV infected cells, there was no increase in cell death (Fig. 4D). Next, we looked at the IFN sensitivity of MPXV with the pancaspase inhibitor. The intermediate IFN sensitivity of MPXV was increased with the addition of zVAD to IFN treated L929 cells. Without zVAD, approximately 50% plaque reduction was observed with 10 U of IFN in the case of MPXV, whereas VACV-E3LΔ37N required only 3 U of IFN. Treatment with zVAD increased IFN sensitivity of MPXV infected cells to resemble VACV-E3LΔ37N with 3 U of IFN giving a 45% plaque reduction. Treatment with zVAD had no effect on IFN sensitivity with VACV-E3LΔ37N (Fig. 4E). Addition of MLN4924 also increased IFN sensitivity of MPXV and VACV-E3LΔ37N infected cells, with 1 U of IFN reducing plaque formation by 50% (Fig. 4F). These results demonstrated that the addition of the pancaspase inhibitor, zVAD, or the neddylation inhibitor, MLN4924, can effectively enhance the IFN sensitivity of MPXV.

## Discussion

Many viruses activate the host type I IFN pathway, leading to the upregulation of interferon stimulated genes (ISGs). These ISGs encode different antiviral pathways that regulate different aspects of the viral lifecycle, such as PKR, OAS, and ZBP1. Viruses, on the other hand, have evolved ways to circumvent these pathways. The VACV E3 protein has been shown to bind dsRNA and sequester it away from PKR and OAS [13]. E3 also sequesters Z-RNA to prevent activation of ZBP1 [29], and is necessary for IFN resistance [13]. The E3 homologue in MPXV is naturally truncated at the N-terminus, but it is still able to inhibit antiviral immune responses to at least some extent [19].

In this manuscript, we show that the addition of IFN⍺ effectively inhibits MPXV viral replication *in vitro* in L929 cells. MPXV is also IFN sensitive in a plaque reduction assay in that increasing IFN doses prevents viral spread to surrounding cells. Addition of 10 U of IFN⍺ is able to decrease plaque formation by 50%. Thus, MPXV is the first unmodified orthopoxvirus characterized to be IFN-sensitive in cells in culture. IFN sensitivity of MPXV is dependent on ZBP1, RIPK3, and MLKL, but not on PKR as knocking out ZBP1, RIPK3, or MLKL fully restored IFN resistance of MPXV, but MPXV remained IFN sensitive in the PKR KOs. It was surprising that the IFN sensitivity was dependent on ZPB1, RIPK3, and MLKL, since MPXV did not induce necroptosis. West Nile virus has been shown to be restricted in neurons in a RIPK3-dependent manner that is independent of cell death and results in chemokine production [37]. Zika virus infection in neurons also leads to ZBP1- and RIPK-dependent inhibition in the absence of necroptosis. ZBP1 and RIPK1/RIPK3 activated a restrictive antiviral state by inducing IRG1 to produce itaconate, which inhibited viral replication [38]. Further MPXV experiments will be needed to determine how the IFN sensitivity is dependent on necroptosis proteins in the absence of cell death.

In addition to not inducing necroptosis, MPXV infection led to delayed MLKL-serine phosphorylation. The delay in MLKL phosphorylation was due to degradation of RIPK3 in MPXV-infected cells. Phosphorylation of MLKL could be at least partially restored either by treating with a pancaspase inhibitor, zVAD to inhibit caspase dependent degradation of RIPK3 or treating with a proteosome inhibitor, MLN4924, to inhibit the vIRD dependent degradation of RIPK3. Despite restoring MLKL-phosphorylation, neither treatment with zVAD nor MLN4924 induced cell death in MPXV infected cells. Treatment with zVAD led to MLKL-phosphorylation, migration of phosphorylated MLKL to membranes, but not trimerization nor cell death. Similarly, treatment with MLN4924 led to increased MLKL phosphorylation but no trimerization. This suggests that MPXV has evolved mechanisms to inhibit necroptosis downstream of MLKL phosphorylation.

While zVAD treatment did not increase induction of necroptotic cell death it did increase sensitivity of MPXV to IFN-treatment. At a concentration of 10 U of IFN⍺, zVAD addition increased MPXV plaque reduction from 50% plaques to 10%. These data show the potential that a pancaspase inhibitor in conjunction with IFN-treatment, or treatment with an IFN-inducer could be used as a MPXV treatment in infected individuals. Future MPXV experiments are planned to test an IFN enhancer with and without a pancaspase inhibitor in an *in vivo* model.

MPXV contains a functional vIRD protein that leads to RIPK3 cleavage and inhibition of necroptosis. We show that RIPK3 expression in MPXV infected cells is greatly reduced compared to VACV and mock infected cells, however it is not completely degraded as in CPXV infected cells. This could be due to a timing issue and at later times post infection, we expect RIPK3 to be completely degraded in MPXV-infected cells. Inhibition of the host SKP1-Cullin1-F-Box (SCF) machinery by the neddylation inhibitor, MLN4924, did not lead to necroptosis as shown by cell viability assay. We observed the same phenomenon as we saw with the addition of the pancaspase inhibitor, zVAD-FMK, in that MLKL gets phosphorylated and travels to the membrane but does not trimerize or cause cell death. MLKL trimerization is the last step in necroptosis required for cell lysis [25], [26]. Thus, even if the host is able to inhibit RIPK3 degradation, MPXV has evolved an additional way to inhibit necroptosis through an unknown mechanism inhibiting MLKL trimerization and cell lysis. Further studies aim to identify the late inhibition of necroptosis downstream of MLKL phosphorylation and characterize the interferon sensitivity in the absence of death.

## Materials and Methods

### Cell lines and treatments

Cells were maintained in 37℃ in a 5% CO_2_ atmosphere. Murine fibroblasts L929 were obtained from ATCC and maintained in Eagle’s Minimum Essential Medium (MEM) (Corning) supplemented with 5% fetal bovine serum (FBS, HyClone). BSC40 cells were maintained in Dulbecco’s Modified-Minimal Essential Medium (DMEM), supplemented with 5% FBS. Pretreatment with mouse IFN⍺ (Calbiochem) was used at 100 U/mL for 1 hour prior to viral infections. To inhibit caspase activity, the pancaspase inhibitor, zVAD-FMK (ApexBio) was used at 50 µM for 1 hour prior to and during infection. For TNF-induced necroptosis, zVAD-FMK was used for pretreatment and TNF-⍺ (Sigma) was added at 20 ng/mL. For RIPK3 degradation inhibition, the neddylation inhibitor MLN 4924 (ApexBio) was added at 1 µM for 1 hour prior to and during infection.

### Viruses

For wtVACV infections, Western Reserve (WR) strain was used. The VACV mutant containing the 37-amino acid N-terminal deletion of E3 (VACV-E3Δ37N), was constructed as previously described [15], [30]. The MPXV strains used in this paper were WR 7-61 (WRAIR) and Zaire 79 (V79-I-005). Non-select agent MPXV experiments were performed in a BSL2+ virus lab and select agent MPXV were performed in BSL3 conditions in accordance with the protocols approved by Arizona State University and the CDC. The CPXV strain used was Brighton red and a kind gift from Grant McFadden’s lab. For all infections, an MOI of 5 was used unless otherwise specified.

### IFN sensitivity plaque assays

L929 and L929 CRISPR/KO cells were seeded in a 12-well tissue culture plate (VWR) to be 95% confluent on the day of infection and treated with increasing doses of IFN⍺ (0-100U/mL) for 18 hours prior to infection. The cells were then infected with 70 PFU of wtVACV, VACV-E3Δ37N, MPXV, or VACV-E3Δ26C and infected for 1 hour with rocking every 10 minutes. After 3-4 days, VACV and VACV mutant infections were stained with crystal violet and plaques counted. For MPXV, plaque formation took 6-7 days, then cells were stained with crystal violet and plaques counted.

### Cell viability assays

L929 cells were plated on a 48-well CytoOne tissue culture-treated plates (USA Scientific) and pretreated with 100 U of IFN⍺ for 18 hours. Cells were infected with the indicated viruses at an MOI of 5 and incubated at 37℃ for 1 hour with rocking every 10 minutes. After infection, cells were overlayed with MEM that contained Hoechst for nuclear staining, and 1 µM SYTOX Green for membrane integrity. The cells were incubated at 37℃ with 5% CO_2_ on an EVOS FL auto imaging microscope (Invitrogen) with an onstage incubator and images were taken every hour for 16 hours.

### Growth kinetics

L929 and CRISPR/Cas9 KO cells were seeded on a 12-well tissue culture plate (VWR) to be 80% on the day of infection. After attachment, cells were pretreated with 100 U of IFN⍺ for 18 hours. Cells were infected with the indicated viruses at an MOI of 5 and incubated at 37℃ with 5% CO_2_ for 1 hour with rocking every 10 minutes. After infection, the cells were washed 3 times with prewarmed media prior to overlay with growth media. The viral samples were harvested at 1, 24, and 48 hours post infection and underwent 3 freeze/thaws. The viral titers were determined by plaquing in BSC40s followed by crystal violet staining.

### Immunofluorescence assay

The LIVE/DEAD Fixable stain kit (ThermoFisher) was used for immunofluorescent (IF) viability assay. L929 cells were seeded to be 90% confluent on the day of infection and pretreated with 100 U of IFN⍺ for 18 hours. Cells were infected with VACV-E3Δ37N or MPXV at an MOI of 5 and infected for 1 hour, followed by growth media overlay. After 16 hours post infection, cells were washed 3X with prewarmed 1X PBS and LIVE/DEAD green fluorescent dye (1:800) was added and incubated at RT for 30 minutes, protected from light. Cells were washed once with 1X PBS and fixed with 4% formaldehyde and incubated at RT for 15 minutes. Once the formaldehyde was removed, 0.125M Glycine was added to the cells to quench any remaining formaldehyde. The cells were washed 3X with 1X PBS and permeabilized by washing 2X with ice-cold PBS. Blocking was done with 3% BSA, 0.3% gelatin, and 0.1% TritonX-100 (TX-100) at RT for 1 hour. The primary antibody (E3 at 1:250) was added in GT-PBS1 (0.3% gelatin, 0.1% TX-100 in PBS) and incubated overnight at 4℃. The cells were washed 2X with GT-PBS1 and the secondary antibody was added (Texas red at 1:250) in GT-PBS1 at RT in the dark for 2 hours. The cells were washed 1X with GT-PBS1 and 1X with PBS and visualized in the EVOS FL auto imaging microscope.

### Protein extraction and Western blot analysis

Cells were seeded in a 12-well tissue culture-treated dish to be 80% confluent on the day of infection. Following viral infection, the cells were scraped, pelleted, washed once with 1X PBS, and lysed in RIPA lysis buffer (150 mM NaCl, 1% P-40, 0.5% sodium deoxycholate, 0.1% SDS, 25 mM Tris pH 7.4) with 1X Halt Protease and Phosphatase Inhibitor Cocktail (Thermo Scientific) and incubated on ice for 5 minutes. For phosphorylated MLKL, QIAshredder columns (Qiagen) were used following cell lysis. Proteins were separated on SDS/Page, transferred to a nitrocellulose membrane with 10 mM 3-(cyclohexylamino)-1-propane sulfonic acid (CAPS), 20% methanol, pH 11 at 100V for 1 hour and 15 minutes, and blocked in 3% bovine serum albumin (BSA). For trimerized MLKL samples, SDS/Page was run under non-reducing conditions and the proteins were transferred at 70V for 3 hours. The following antibodies were used: ZBP1 (Zippy-1, Adipogen), RIPK3 (ProSci), pMLKL (abcam), MLKL (Cell Signaling Technology), β-Actin (Santa Cruz Biotechnology), and PKR (Santa Cruz Biotechnology). Secondary antibodies were goat anti-rabbit IgG or goat anti-mouse IgG (Cell Signaling Technology), depending on the primary antibody used. The proteins were visualized by chemiluminescence with either SuperSignal West Pico PLUS or Dura substrate (ThermoFisher).

### MLKL localization with TX-114

Cells were seeded on 6-well tissue culture-treated plates to be 80% confluent on the day of infection and pretreated with 100 U IFN⍺. After infection, the cells were scraped, pelleted, and washed with cold 1X PBS, and lysis buffer was added (20mM HEPES, 150 mM NaCl, 2% Triton X-114) and incubated on ice for 30 minutes. The samples were then centrifuged at 15,000 x g for 5 minutes at 4℃, the supernatant was harvested and incubated at 37℃ for 5 minutes, followed by centrifugation at 1,500 x g for 5 minutes at room temperature. The top aqueous layer was separated, additional TX-114 lysis buffer was added and the samples were incubated at 37℃ for additional 3 minutes, followed by centrifugation at 1,500 x g for 5 minutes to remove any contamination from the detergent layer. The samples were saved as Aq samples. The detergent enriched layer was diluted with basal buffer (20 mM HEPES, 150 mM NaCl) to the same volume as the detergent soluble fraction, incubated at 4℃ for 15 minutes, incubated at 37℃ for 3 minutes and centrifuged at 1,500 x g for 5 minutes. The samples were diluted with basal buffer to the same volume as the Aq samples and saved as Det. The proteins were then visualized by Western blot as described above.

### L929 CRISPR KO cell lines

L929 cells were transduced with Edit-R Lentiviral particles containing Cas9 sgRNA (Horizon Discovery) according to manufacturer’s instruction and expression of Cas9 was verified by Western blot. The Cas9 expressing L929 cells were then transduced to KO the following proteins: ZBP1, RIPK3, PKR, and MLKL. The cells were infected with Edit-R Lentiviral specific sgRNAs (Horizon Discovery) at an MOI of 0.3 and the remaining protocol was followed according to the manufacturer’s instruction. The protein KOs were confirmed via Western blot.

## Notes

### Competing Interest Statement

The authors have declared no competing interest.

## References

[1] A. M. Likos et al., “A tale of two clades: monkeypox viruses,” Journal of General Virology, vol. 86, no. 10, pp. 2661–2672, doi: 10.1099/vir.0.81215-0.

[2] A. W. Rimoin et al., “Major increase in human monkeypox incidence 30 years after smallpox vaccination campaigns cease in the Democratic Republic of Congo,” PNAS, vol. 107, no. 37, pp. 16262–16267, Sep. 2010, doi: 10.1073/pnas.1005769107.

[3] “The Global Eradication of Smallpox: Final Report of the Global Commision for the Certification of Smallpox Eradication: World Health Organization.” 1980. [Online]. Available: http://apps.who.int/iris/bitstream/handle/10665/39253/a41438.pdf?sequence=1&isAllowed=y

[4] K. D. Reed et al., “The detection of monkeypox in humans in the Western Hemisphere,” New England Journal of Medicine, vol. 350, no. 4, pp. 342–350, 2004.

[5] N. Erez et al., “Diagnosis of Imported Monkeypox, Israel, 2018 - Volume 25, Number 5—May 2019 - Emerging Infectious Diseases journal - CDC”, doi: 10.3201/eid2505.190076.

[6] A. Vaughan et al., “Human-to-Human Transmission of Monkeypox Virus, United Kingdom, October 2018 - Volume 26, Number 4—April 2020 - Emerging Infectious Diseases journal - CDC”, doi: 10.3201/eid2604.191164.

[7] S. E. F. Yong et al., “Imported Monkeypox, Singapore - Volume 26, Number 8— August 2020 - Emerging Infectious Diseases journal - CDC”, doi: 10.3201/eid2608.191387.

[8] A. K. Rao, “Monkeypox in a Traveler Returning from Nigeria — Dallas, Texas, July 2021,” MMWR Morb Mortal Wkly Rep, vol. 71, 2022, doi: 10.15585/mmwr.mm7114a1.

[9] “Multi-country outbreak of monkeypox, External situation report #1 - 6 July 2022.” Accessed: Jul. 08, 2022. [Online]. Available: https://www.who.int/publications/m/item/multi-country-outbreak-of-monkeypox--external-situation-report--16-july-2022

[10] S. Reardon, “Mpox is spreading rapidly. Here are the questions researchers are racing to answer,” Nature, vol. 633, no. 8028, pp. 16–17, Aug. 2024, doi: 10.1038/d41586-024-02793-9.

[11] J. G. Rizk, G. Lippi, B. M. Henry, D. N. Forthal, and Y. Rizk, “Prevention and Treatment of Monkeypox,” Drugs, vol. 82, no. 9, pp. 957–963, Jun. 2022, doi: 10.1007/s40265-022-01742-y.

[12] G. Yang et al., “An Orally Bioavailable Antipoxvirus Compound (ST-246) Inhibits Extracellular Virus Formation and Protects Mice from Lethal Orthopoxvirus Challenge,” J Virol, vol. 79, no. 20, pp. 13139–13149, Oct. 2005, doi: 10.1128/JVI.79.20.13139-13149.2005.

[13] H.-W. Chang, J. C. Watson, and B. L. Jacobs, “The E3L gene of vaccinia virus encodes an inhibitor of the interferon-induced, double-stranded RNA-dependent protein kinase,” Proceedings of the National Academy of Sciences, vol. 89, no. 11, pp. 4825–4829, 1992.

[14] H.-W. Chang and B. L. Jacobs, “Identification of a conserved motif that is necessary for binding of the vaccinia virus E3L gene products to double-stranded RNA,” Virology, vol. 194, no. 2, pp. 537–547, 1993.

[15] H.-W. Chang, L. H. Uribe, and B. L. Jacobs, “Rescue of vaccinia virus lacking the E3L gene by mutants of E3L.,” Journal of virology, vol. 69, no. 10, pp. 6605–6608, 1995.

[16] H. Yuwen, J. H. Cox, J. W. Yewdell, J. R. Bennink, and B. Moss, “Nuclear localization of a double-stranded RNA-binding protein encoded by the vaccinia virus E3L gene,” Virology, vol. 195, no. 2, pp. 732–744, 1993.

[17] J. C. Watson, H.-W. Chang, and B. L. Jacobs, “Characterization of a vaccinia virus-encoded double-stranded RNA-binding protein that may be involved in inhibition of the double-stranded RNA-dependent protein kinase,” Virology, vol. 185, no. 1, pp. 206–216, 1991.

[18] T. A. Brandt and B. L. Jacobs, “Both carboxy-and amino-terminal domains of the vaccinia virus interferon resistance gene, E3L, are required for pathogenesis in a mouse model,” Journal of virology, vol. 75, no. 2, pp. 850–856, 2001.

[19] W. D. Arndt et al., “Evasion of the Innate Immune Type I Interferon System by Monkeypox Virus,” Journal of Virology, vol. 89, no. 20, pp. 10489–10499, Oct. 2015, doi: 10.1128/JVI.00304-15.

[20] Y. Lin, A. Devin, Y. Rodriguez, and Z. Liu, “Cleavage of the death domain kinase RIP by Caspase-8 prompts TNF-induced apoptosis,” Genes Dev., vol. 13, no. 19, pp. 2514–2526, Oct. 1999.

[21] Y. Cho et al., “Phosphorylation-Driven Assembly of the RIP1-RIP3 Complex Regulates Programmed Necrosis and Virus-Induced Inflammation,” Cell, vol. 137, no. 6, pp. 1112–1123, Jun. 2009, doi: 10.1016/j.cell.2009.05.037.

[22] L. Sun et al., “Mixed Lineage Kinase Domain-like Protein Mediates Necrosis Signaling Downstream of RIP3 Kinase,” Cell, vol. 148, no. 1, pp. 213–227, Jan. 2012, doi: 10.1016/j.cell.2011.11.031.

[23] Z. Cai et al., “Plasma membrane translocation of trimerized MLKL protein is required for TNF-induced necroptosis,” Nature Cell Biology, vol. 16, no. 1, pp. 55– 65, Jan. 2014, doi: 10.1038/ncb2883.

[24] E. S. Mocarski, J. W. Upton, and W. J. Kaiser, “Viral infection and the evolution of caspase 8-regulated apoptotic and necrotic death pathways,” Nature Reviews Immunology, vol. 12, no. 2, pp. 79–88, Feb. 2012, doi: 10.1038/nri3131.

[25] Y. Dondelinger et al., “MLKL compromises plasma membrane integrity by binding to phosphatidylinositol phosphates,” Cell reports, vol. 7, no. 4, pp. 971–981, 2014.

[26] H. Wang et al., “Mixed Lineage Kinase Domain-like Protein MLKL Causes Necrotic Membrane Disruption upon Phosphorylation by RIP3,” Molecular Cell, vol. 54, no. 1, pp. 133–146, Apr. 2014, doi: 10.1016/j.molcel.2014.03.003.

[27] J. W. Upton, W. J. Kaiser, and E. S. Mocarski, “DAI/ZBP1/DLM-1 Complexes with RIP3 to Mediate Virus-Induced Programmed Necrosis that Is Targeted by Murine Cytomegalovirus vIRA,” Cell Host & Microbe, vol. 11, no. 3, pp. 290–297, Mar. 2012, doi: 10.1016/j.chom.2012.01.016.

[28] H. Koehler et al., “Inhibition of DAI-dependent necroptosis by the Z-DNA binding domain of the vaccinia virus innate immune evasion protein, E3,” Proceedings of the National Academy of Sciences, vol. 114, no. 43, pp. 11506–11511, 2017.

[29] H. Koehler et al., “Vaccinia virus E3 prevents sensing of Z-RNA to block ZBP1-dependent necroptosis,” Cell Host & Microbe, vol. 29, no. 8, pp. 1266–1276.e5, Aug. 2021, doi: 10.1016/j.chom.2021.05.009.

[30] T. Shors, K. V. Kibler, K. B. Perkins, R. Seidler-Wulff, M. P. Banaszak, and B. L. Jacobs, “Complementation of Vaccinia Virus Deleted of the E3L Gene by Mutants of E3L,” Virology, vol. 239, no. 2, pp. 269–276, Dec. 1997, doi: 10.1006/viro.1997.8881.

[31] S. Feng et al., “Cleavage of RIP3 inactivates its caspase-independent apoptosis pathway by removal of kinase domain,” Cellular Signalling, vol. 19, no. 10, pp. 2056–2067, Oct. 2007, doi: 10.1016/j.cellsig.2007.05.016.

[32] S. Alvarez-Diaz et al., “The Pseudokinase MLKL and the Kinase RIPK3 Have Distinct Roles in Autoimmune Disease Caused by Loss of Death-Receptor-Induced Apoptosis,” Immunity, vol. 45, no. 3, pp. 513–526, Sep. 2016, doi: 10.1016/j.immuni.2016.07.016.

[33] W. J. Kaiser et al., “RIP3 mediates the embryonic lethality of caspase-8-deficient mice,” Nature, vol. 471, no. 7338, pp. 368–372, Mar. 2011, doi: 10.1038/nature09857.

[34] A. Oberst et al., “Catalytic activity of the caspase-8–FLIPL complex inhibits RIPK3-dependent necrosis,” Nature, vol. 471, no. 7338, pp. 363–367, 2011.

[35] Z. Liu et al., “A class of viral inducer of degradation of the necroptosis adaptor RIPK3 regulates virus-induced inflammation,” Immunity, vol. 54, no. 2, pp. 247–258.e7, Feb. 2021, doi: 10.1016/j.immuni.2020.11.020.

[36] Y. Taguchi and H. M. Schätzl, “Small-scale Triton X-114 Extraction of Hydrophobic Proteins,” Bio Protoc, vol. 4, no. 11, p. e1139, Jun. 2014, doi: 10.21769/BioProtoc.1139.

[37] B. P. Daniels et al., “RIPK3 Restricts Viral Pathogenesis via Cell Death-Independent Neuroinflammation,” Cell, vol. 169, no. 2, pp. 301–313.e11, Apr. 2017, doi: 10.1016/j.cell.2017.03.011.

[38] B. P. Daniels et al., “The Nucleotide Sensor ZBP1 and Kinase RIPK3 Induce the Enzyme IRG1 to Promote an Antiviral Metabolic State in Neurons,” Immunity, vol. 50, no. 1, pp. 64–76.e4, Jan. 2019, doi: 10.1016/j.immuni.2018.11.017.

